# Single-time-point shotgun metagenomics of four Irish Integrated Constructed Wetlands reveals microbial dynamics in wastewater treatment

**DOI:** 10.64898/2026.09.08.750087

**Authors:** Anna Tumeo, Gaia Streparola, Caolan Harrington, Aila Carty, Finola Leonard, Catherine Burgess, Dearbháile Morris, Georgios Miliotis

## Abstract

Over the past 20 years, the Integrated Constructed Wetland (ICW) concept has been applied in Ireland for wastewater treatment, offering a nature-based solution for reducing pollutants and antimicrobial resistance genes (ARGs) in effluents and receiving environments. However, wastewater microbial communities remain inadequately characterized and their dynamics across treatment largely unexplored. Here, we present a culture-independent investigation of four Irish ICWs, aiming to advance our understanding of microbial dynamics in wastewater treatment. Shotgun metagenomic sequencing was conducted on influent and effluent samples collected in biological triplicates from four ICWs treating agricultural, companion animal, industrial, and municipal wastewater, alongside eight positive and ten negative controls for filtering and sequencing. End-to-end metagenomic analysis was performed with SqueezeMeta. Taxonomic placement of high-quality metagenome-assembled-genomes (HQ-MAGs) was confirmed with GTDB-Tk. Decontamination and statistical analyses were conducted in R. Coassembled contigs were screened for ARGs, virulence- and plasmid-associated sequences. Results revealed statistically significant shifts in taxonomic and functional profiles across treatment. Effluent populations exhibited generally higher richness than corresponding influents, yet were more similar to one another across locations. Multi-log-scale increases were observed in the relative abundance of environmental genera (*Legionella, Methylotenera, Thiothrix*), alongside reductions in common faecal/human indicators (*Bacteroides, Lactococcus, Prevotella*), and up to 80% ARGs removal. Collectively, our findings provide insights into ICW efficacy, showing that they can drive marked shifts in wastewater microbiome and ARG reduction. Our culture-independent approach retrieved 34 HQ-MAGs, including 19 potentially novel taxa, uncovering previously uncharacterized microbial diversity that is potentially unique to the Irish environment and warrants further investigation.

## 1. Introduction

Wastewater is an environment rich in pollutants and nutrients that fosters complex and dynamic microbial communities. Research to date on human and animal wastewater from urban and industrial sources reports the consensus identification of Proteobacteria (Pseudomonadota), Actinobacteria (Actinomycetota), Firmicutes (Bacillota), and Bacteroidetes (Bacteroidota) as dominant taxa, likely owing to their versatile metabolic roles in nutrient cycling and contaminant degradation, as well as to their ability to tolerate environmental stresses and degrade complex organic compounds [1–4]. The composition of these communities at deeper taxonomic levels varies considerably depending on wastewater type, local environment, and characteristics and extent of anthropogenic contamination.

Of particular concern, growing evidence implicates wastewaters as environmental reservoirs for antimicrobial resistance (AMR) and documents the occurrence of opportunistic pathogenic taxa of primary public health interest including *Acinetobacter, Enterobacter, Klebsiella*, and *Pseudomonas* [5–8]. As human, agricultural, and industrial activities can amplify dispersion of these microbial contaminants into the surrounding environment [9], wastewater treatment is critical to prevent their transmission to humans, animals, and the food chain.

Integrated constructed wetlands (ICWs) are an innovative Irish solution to wastewater treatment that mimic natural wetlands to degrade, adsorb, and transform pollutants by using vegetation, soil, and associated microbial communities [10]. In ICWs, autochthonous bacterial populations may play a role in the control of microbial contaminants through, for example, the production of microbial exudates and competition for limiting nutrients [11]. By combining ecological principles with engineering design, ICWs show promise as sustainable and cost-effective alternatives to conventional wastewater treatment. However, little is known of the microbiome that may constitute the basis of their mechanism of action, which restricts our capacity to optimize treatment strategies [12]. Treatment-induced changes in influent wastewater microbial populations, including removal of pathogenic taxa and involvement of microbial communities in nutrient cycling, have also been minimally explored. This is due, at least in part, to the novelty of ICWs as wastewater treatment systems, which have not yet been extensively characterized. In addition, narrow-target methods, including culture-based assays and targeted quantitative PCR, cannot resolve the full breadth of microbial diversity and functional potential that may contribute to ICW performance.

Consequently, key research questions remain inadequately addressed, including the dynamics of influent and autochthonous microbial communities in ICWs and the extent of AMR and opportunistic pathogens mitigation that these systems can achieve.

Here, we use shotgun metagenomics to compare influent and effluent microbial communities sampled during a single-time-point campaign at four Irish ICWs treating contrasting wastewater sources. Through the reconstruction of influent and effluent bacterial populations and their changes upon treatment, our findings provide insights into the efficacy of these nature-based wastewater treatment systems at mitigating AMR and pathogenic taxa. Furthermore, our findings show preliminary evidence of post-treatment shifts in microbial functions related to nitrogen, phosphorus, and sulfur transformation, suggesting that ICWs may select for communities with altered nutrient-acquisition and transformation potential. By incorporating independent biological replicates and multiple positive and negative controls alongside state-of-the-art bioinformatic pipelines, this work additionally aims to contribute to the development of more robust, contamination-aware wastewater microbiome metagenomic approaches for evaluating microbial contaminant removal in wastewater treatment systems.

## 2. Methods

### 2.1 Experimental design and site description

Four ICWs spanning different wastewater types, namely agricultural (AG), companion animal (CA), industrial (IND), and municipal (MUN), were selected across Ireland. AG wastewaters include stormwater runoff from the yard and roof of a dry stock farm. The associated ICW covers a treatment area of 0.50ha and consists of three cells, the first being adjacent to the yard and the latter two to the receiving stream. CA wastewaters include washdown and foul waters from an animal shelter and veterinary practice facility. The associated ICW was developed in 2018 and provides treatment through four cells over a treatment area of 0.50ha, and discharge waters enter a receiving stream. IND wastewaters are secondary-treated wastewaters from a large food distribution facility, fed to the ICW before discharge to a receiving stream. The ICW comprises one settlement cell for removal of heavy solids and a series of four cells over a treatment area of 0.85ha. The MUN ICW was developed in 2021 as part of a stormwater misconnection project aiming to improve surface water quality in receiving waters. The ICW consists of two cells and a treatment area of 0.50ha.

Sampling was conducted in December 2024. Samples (V=1 L) were collected from both influent (pre-treatment) and effluent (post-treatment) wastewater in independent biological triplicates using sterile collection bottles and stored on ice until processing. Environmental (temperature, rainfall, relative humidity, wind speed), water quality (sample temperature and pH), and wetland operational parameters were recorded at the time of sampling to ensure capture of all relevant metadata (**Table S1**).

### 2.2 Sample processing including positive and negative controls

Environmental samples (n=24) were filtered through 0.45 µm pore-sized, sterile, cellulose nitrate membrane Microsart® filters (Sartorius, Goettingen, Germany) up to filter saturation using a vacuum manifold under controlled pressure. Filters were stabilized in DNA/RNA Shield (Zymo Research, USA) and stored at room temperature until DNA extraction.

Eight positive filtration controls were incorporated in the study and processed alongside environmental samples to assess filtration efficacy. These consisted of equal aliquots (10 µL for influent and 5 µL for effluent samples) of reconstituted ATCC® MSA-2003™ spiked into one replicate (V=200 mL) of each sample type. MSA-2003™ is an even mix of whole cell material from ten authenticated ATCC Genuine Cultures®, used as a mock microbial community to mimic a mixed metagenomic sample.

Six types (i-vi) of negative controls were incorporated to detect the introduction of contamination and minimize bias at all stages of the analysis, including: three negative controls for filtration, consisting of (i) pure sterile PBS (V=200 mL) filtered alongside the environmental samples and (ii, iii) three filters each from two separate production batches (251100267, 252800617), suspended in DNA/RNA Shield (1.5 mL) without additional sample DNA; three “kitome” controls used to assess the presence of any background contamination associated with the employed kits and reagents, including (iv) pure DNA/RNA Shield (200 µL), (v) pure sterile PBS (5 mL), and (vi) one DNA extraction/sequencing negative control as DNA-free water (200 µL) subjected to DNA extraction and library preparation alongside the environmental samples. All controls (n=10) were processed simultaneously with the rest of the samples to ensure equivalent handling and storage conditions. DNA was extracted from the stabilized environmental membrane filters and corresponding control materials by BGI Genomics using the MGIEasy Stool Microbiome DNA Extraction Kit (MGI Tech Co., Ltd., Shenzhen, China) according to the manufacturer’s instructions. Sequencing libraries were prepared using the MGIEasy Fast FS DNA Library Prep Set (Cat. No. 940-000029-00, MGI Tech) and sequenced on a DNBSEQ-T7 platform using 150-bp paired-end reads. Target sequencing outputs were 15 Gb per environmental sample and 5 Gb per control. Environmental samples and controls were processed using the same DNA-extraction and library-preparation workflow.

### 2.3 Bioinformatic analysis

Quality-checked reads (Q score ≥ 33) were given as input to SqueezeMeta v1.6.3 [13] for assembly, gene prediction, taxonomic assignment, functional annotation, and binning. For environmental samples, replicates (n=3) of the same type were coassembled to minimize the percentage of non-assembled reads while maximizing binning capacity. The R package Phyloseq was used to handle raw taxonomic assignment data, excluding taxonomic assignments below a minimum threshold of 5 reads. Bacterial taxonomic and functional features potentially associated with contamination were identified with the R package decontam [14], using the prevalence method and a threshold of 0.6. For decontamination, raw counts of taxonomic assignments were normalized across environmental samples and controls to account for differences in sequencing depth. Alpha and beta diversity analyses were performed using the R package vegan v2.7-1. Statistical comparison of alpha diversity indices between influent and effluent unpaired groups was performed with a t-test. The Bray-Curtis dissimilarity metric was used to assess beta diversity and functional diversity, which were visualized through Non-Metric MultiDimensional Scaling (NMDS) ordination plots. The R package Microbiome Regression-Based Kernel Association Test (MiRKAT) v1.2.1 [15] was used to determine statistical significance of clustering between influent and effluent microbial communities, and PERMANOVA for clustering between environmental samples and negative controls. PERMDISP was used to determine homogeneity within groups. For all tests, statistical significance was defined as p < 0.05. Changes in relative and absolute counts of taxonomic and functional features were expressed as log2 fold-change (log2FC), calculated from the mean abundance values across triplicate influent and effluent samples:

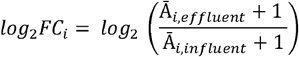

with a pseudo-count of 1 added to avoid undefined values where a feature was absent from one sample group.

Site-specific coassembled contigs were screened for ARGs, virulence factors (VFs), and plasmid sequences using Abricate v1.0.1 against CARD v4.0.1 [16], VFDB 2025 [17], ecOH [18], and plasmidfinder v2.1 [19] databases. Hits with >60% coverage and >80% identity were linked to the taxonomic assignment of the corresponding contigs using a custom script. Taxonomic placement of high-quality (≥95% completeness; ≤5% contamination) metagenome-assembled genomes (MAGs) identified and generated by SqueezeMeta was verified using GTDB-Tk v2.6.1 with database r226 [20].

## 3. Results

### 3.1 Microbial load across environmental samples and negative controls

Domain- and genus-specific normalized read counts are available as **Table S2: Domain; Genus**. Shotgun metagenomic sequencing of all environmental samples (including replicates, n=24) yielded a total of 2.9 x 10^9^ reads (median: 120,122,102). Bacterial reads constituted between 38 and 94% (median: 52%) of the total reads per sample, adding up to a total of 1.6 x 10^9^ (median: 5.8 x 10^7^) bacterial reads obtained across all 24 environmental samples. Retrieval of the ten taxa represented in ATCC® MSA-2003™ mock community through sequencing of the positive controls is shown in **Table S2: Positive Controls**. Collectively, negative controls (n=10) yielded 4.8 x 10^6^ reads (pre-normalization), with >99.9% of the signal originating from the negative controls for filtration (**Table S2: Negative Controls**). Over 2.6 x 10^6^ bacterial reads assigned to 631 bacterial taxa were detected in the negative controls. Among these, ten bacterial taxa were flagged as contaminants upon control-informed data filtering, therefore removed from the dataset and excluded from downstream analyses (**Table S3**).

A total of 34 HQ-MAGs were retrieved from the environmental samples through metagenome binning (**Table S4**). Among these, only 15 were taxonomically classified at the species level.

### 3.2 Alpha diversity

Alpha diversity analysis based on shotgun metagenomic sequencing data showed significant differences in microbial diversity before and after treatment (**Figure 1**). In particular, significantly (t-test: p ≤ 0.01) higher counts of observed genera were consistently observed in the effluents of all four sampled ICWs compared to the corresponding influents (**Figure 1A**). Shannon and Inverse Simpson indices (**Figure 1B-C**) were also higher for the effluents of the agricultural, companion animal, and industrial ICWs compared to their influents, although divergence between Inverse Simpson values was not statistically significant (t-test: p=0.1) for the agricultural ICW. In the case of the municipal ICW, influent and effluent replicates yielded comparable alpha diversity according to both Shannon and Inverse Simpson indices.

**Figure 1:**
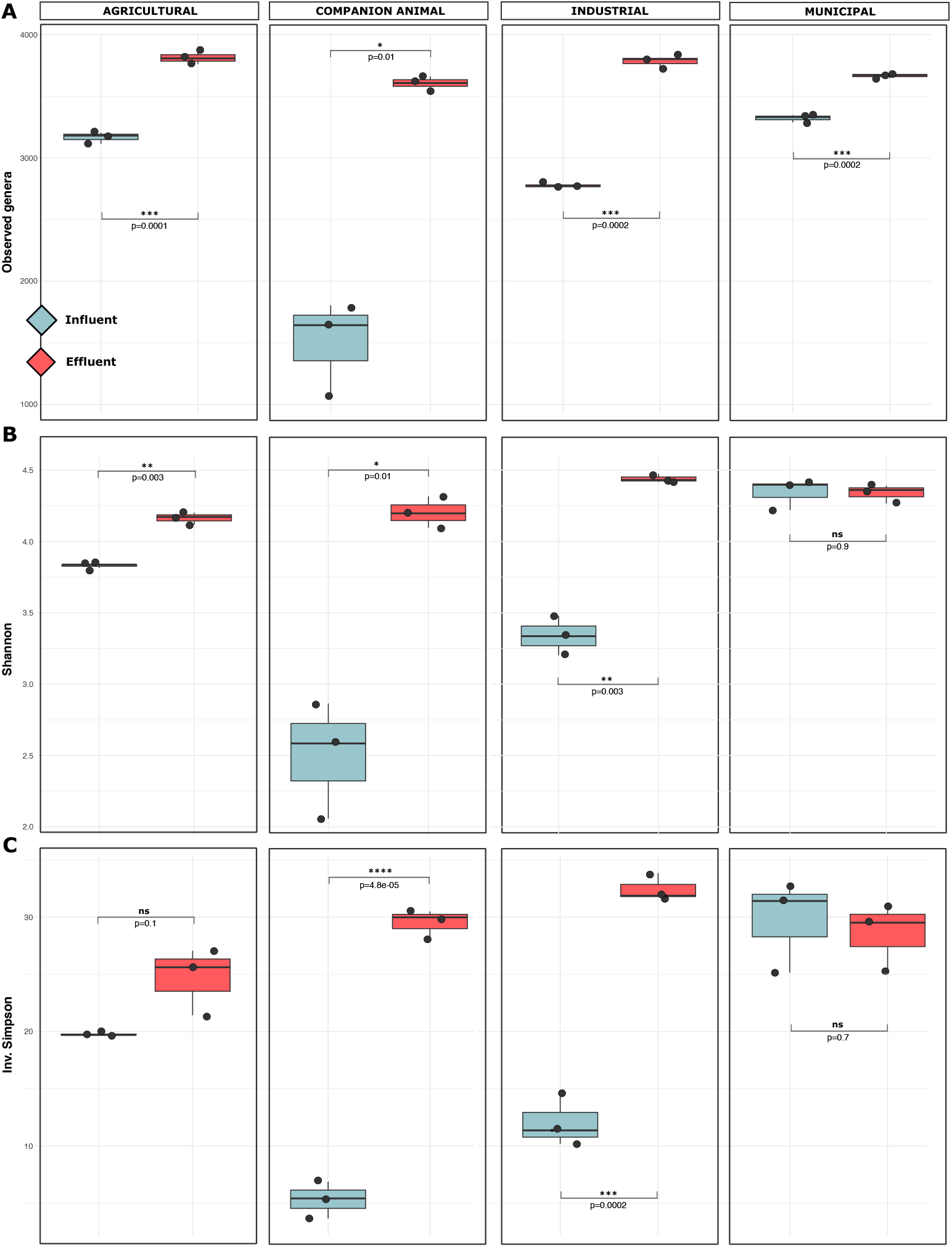
Alpha diversity. Shifts in alpha diversity between influent (cyan) and effluent (red) samples. For each sampled ICW, alpha diversity was calculated as (**A**) counts of observed genera, (**B**) Shannon index, and (**C**) Inverse Simpson index. Values are reported for all biological replicates (n=3). Statistical significance was inferred with a t-test.

### 3.3 Beta diversity

Beta diversity analysis using the Bray-Curtis dissimilarity metric showed site-specific clustering of the environmental samples, indicating the presence of highly distinct communities in the four sampled ICWs (**Figure 2A**). This is supported by tight clustering between triplicates of the same sample type, indicating biological and technical consistency. For each ICW, NMDS showed separation between influent and effluent communities, suggesting treatment-induced shifts in their microbial composition, with effluent communities from different sites often clustering more closely together than with their corresponding influents. True samples formed a cluster separated from negative controls (PERMANOVA: R^2^ = 0.16; p = 0.003, PERMDISP: F = 2.11; p = 0.156), suggesting limited contribution of reagent contamination to the observed microbial profiles. However, negative controls for filtration yielded >1.0 ×10^6^ reads, indicating a substantial amount of background contamination in the hardware used for processing the sample (**Figure 2B**). Community profiling based on shotgun metagenomic sequencing data uncovered major shifts in the microbial composition of influent and effluent samples of each ICW (**Figure 2C**). Dominant influent taxa included common gut indicators such as *Prevotella, Bacteroides, Lactococcus, Clostridium*, and environmental bacteria such as *Acinetobacter, Malikia*, and *Shewanella*, while genera normally associated with sludge and groundwater including *Methylotenera, Thiothrix*, and *Legionella* were predominantly found in the effluents. However, microbial profiling uncovered some of the most abundant taxa in the influent persisting as dominant after treatment. These included *Aliarcobacter* and *Pseudomonas* in the companion animal ICW, and *Flavobacterium* in the industrial ICW. In addition, the municipal ICW showed limited changes in the taxa dominating influent and effluent microbial communities.

**Figure 2:**
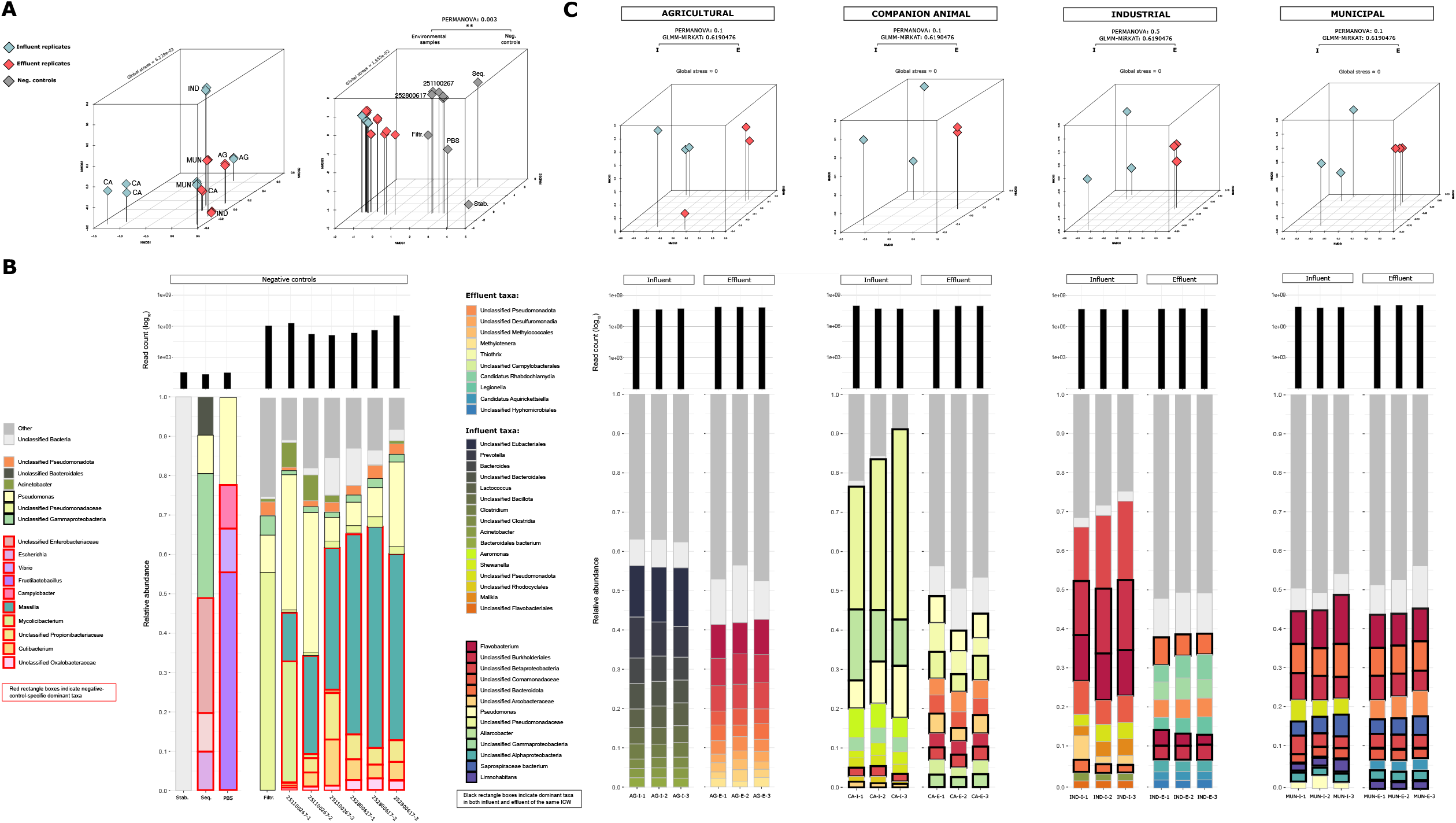
Beta diversity and microbial composition. (**A**) NMDS ordination plots showing clustering by compositional similarity of the environmental samples (including replicates, n=24) respectively (left) with and (right) without negative controls (n=10). In the figure, “Stab.” is the stabilization reagent negative control; “Seq.” is the negative control for library preparation/sequencing; “PBS” is pure, unprocessed PBS; “Filtr.” is pure PBS filtered through the same cellulose nitrate filters as the environmental samples; 251100267 and 252800617 refer to two production batches of cellulose nitrate membrane Microsart® filters. Three filters from each batch were used as negative controls; (**B**) Predominant microbial taxa identified in filtration and kitome negative controls; (**C**) Predominant microbial taxa identified in each sampled ICW before and after treatment. For each ICW, NMDS plots signify compositional distinction between influent (cyan) and effluent (red) communities.

### 3.4 Multi-log-scale community shifts across treatment

Figure 3. highlights the bacterial groups showing the greatest statistically significant (p < 0.05) change in relative abundance across treatment at each ICW. The largest post-treatment decreases were dominated by human- and enteric-associated genera, including taxa that contain opportunistic pathogens. Among these, *Avrilella* at the agricultural site, *Megamonas, Morganella*, and *Laribacter* at the companion animal site, and Lelliottia at the industrial site. Unclassified members of *Aeromonadaceae, Fusobacteriaceae*, and *Ignatzschineria*ceae families, which include opportunistic genera such as *Aeromonas* spp. and *Ignatzschineria*, also showed strong reductions at the industrial and companion animal sites, respectively. Across all sites, the largest post-treatment increases were observed for bacterial groups commonly reported in freshwater and wastewater (*Lacunisphaera, Polymorphobacter, Aurantimicrobium, Aquisediminimonas*) and soil (*Archangium, Verrucomicrobium, Ktedonobacter*). Several of the enriched taxa are associated with redox and nutrient cycling (*Thiothrix, Desulfomonile, Sorangineae, Pseudorhodoplanes*), and are anaerobic community members linked to syntrophy (e.g., Syntrophorhabdaceae) or fermentation (Paludibacteraceae). Fewer statistically significant shifts were observed at the municipal site, indicating an overall more stable microbial population throughout treatment. In this case, however, *Ignatzschineria* was found to increase the effluent substantially.

**Figure 3:**
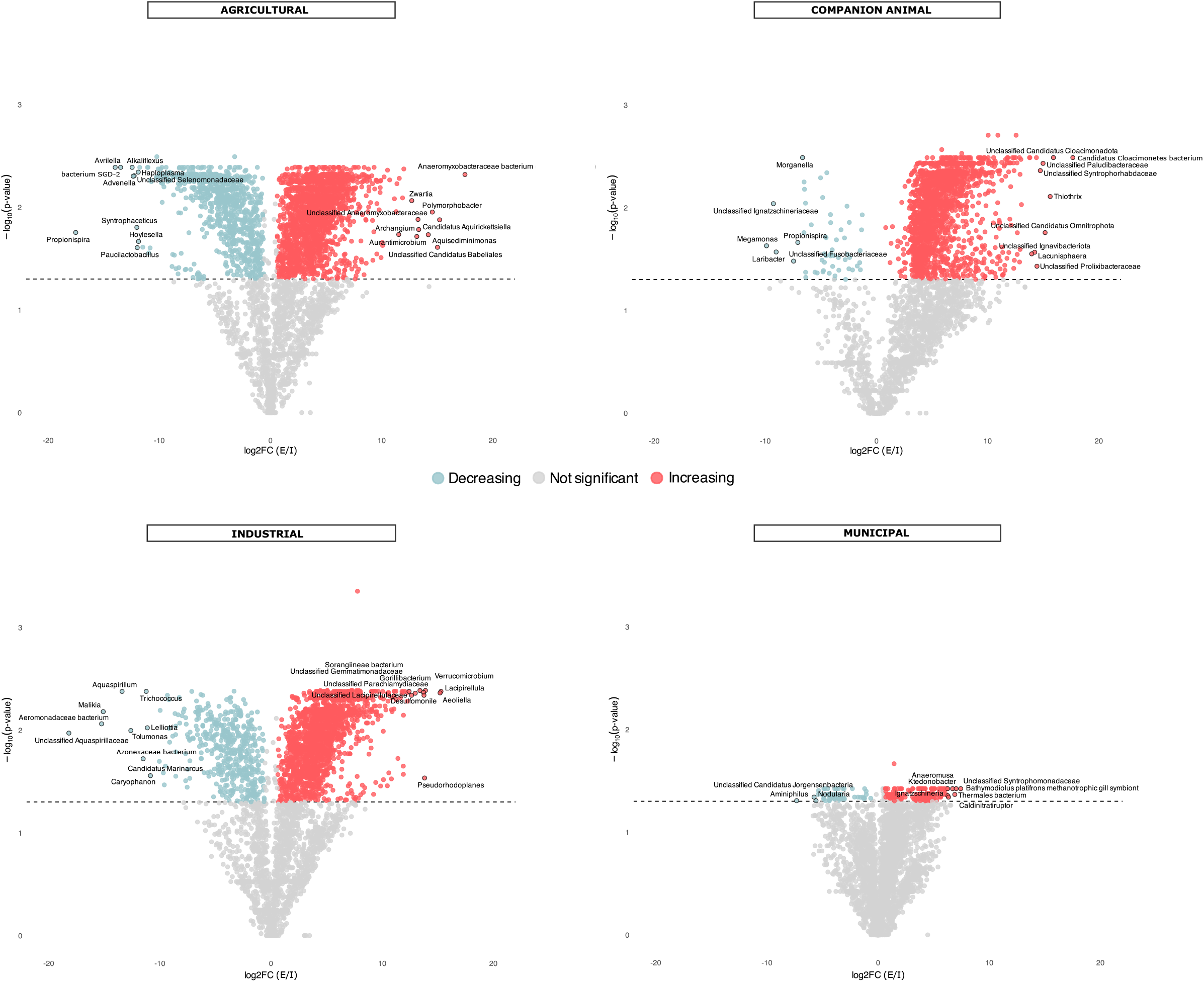
Multi-log-scale community shifts across treatment. Bacterial taxa showing statistically significant (p < 0.05) decreases (cyan) or increases (red) in relative abundance following treatment at each ICW. Changes in the relative abundance of taxonomic features were expressed as log2 fold-change (log2FC), calculated from the mean relative abundance values across triplicate influent and effluent samples.

### 3.5 Functional characterization of influent and effluent microbial communities

Functional assignment of metagenomic contigs yielded a total of 94676 distinct COG/ENOG IDs across all environmental samples (including replicates, n=24) and negative controls (n=10) (**Table S5: Overview**), with ~80% (n=74910) mapped to unknown functions. NMDS showed distinction between influent and effluent microbial communities based on their functional profiles, supported by close clustering of triplicates and separation from the negative control (PERMANOVA: R^2^ = 0.17; p = 0.042) (**Figure 4A**). NMDS confirmed influent- and effluent-specific sample clustering for each site, although limited group size (n=3) hinders statistical significance (**Figure 4B**). Analysis of the differential abundance of annotated functions across influent and effluent of each site revealed that statistically significant changes occurred upon treatment at the agricultural and companion animal sites, while no significant changes occurred at the industrial and municipal sites. Functions that significantly (BH-adjusted p < 0.05) decreased in the treated agricultural effluent included carbohydrate transport and metabolism (e.g., ENOG410ZIAQ, ENOG4111MCP) and transcription (e.g., COG4109, COG4463, ENOG410ZREP, ENOG41124NK). Functions that significantly increased in the treated agricultural effluent at this site were associated with post-translational modification and protein turnover (e.g., COG5647, ENOG410XNV7, ENOG410XP8T, ENOG410XP9Y, ENOG410XSAG) and signal transduction (e.g., ENOG4110KJ7, ENOG410YG82, ENOG41101WB, COG5040, ENOG4112C51, ENOG410XRI7, ENOG41112FU, ENOG410XNRB). Similarly, in the companion animal effluent, significantly decreased functions were associated with carbohydrate transport and metabolism (e.g., ENOG410ZVSA, ENOG410ZVX0, ENOG410YA3A, ENOG41112TP) and transcription (e.g., ENOG411247E, ENOG4111MMN, ENOG4111GHD). Functions that instead significantly increased in the effluent included post-translational modification and protein turnover (e.g., COG5077) and energy production and conversion (e.g., ENOG4110RRB, ENOG4112C99, ENOG4111TBQ). No significant changes were observed in the functional profile of the microbial communities following treatment at the industrial and municipal sites.

**Figure 4:**
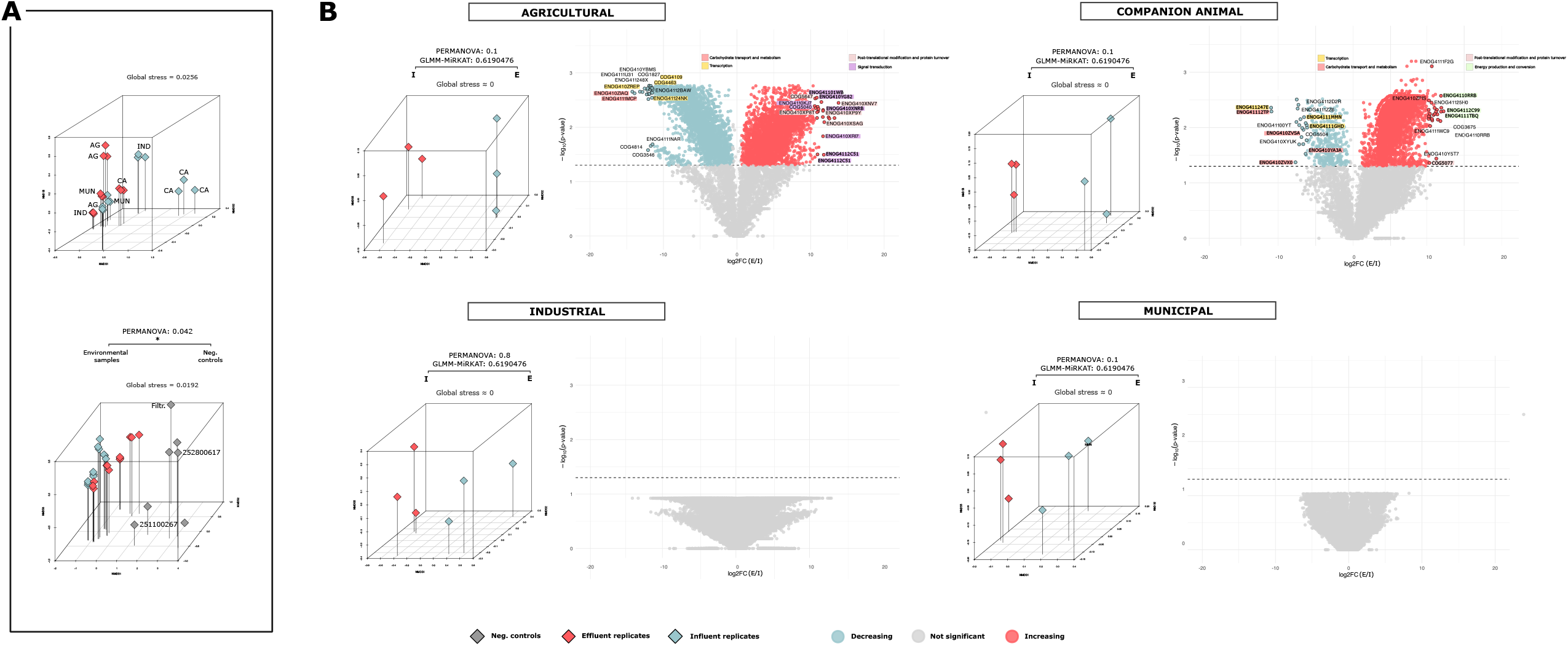
Functional characterization of microbial communities. (**A**) NMDS plots showing separation between environmental samples (including replicates, n=24) based on their functional profiles, respectively (top) excluding and (bottom) including negative controls. Only negative controls with non-zero functional assignments are shown; (B) Microbial functions undergoing statistically significant (BH-adjusted p < 0.05) decreases (cyan) or increases (red) in raw copy abundance following treatment at each ICW. Changes in the absolute abundance of functional features were expressed as log2 fold-change (log2FC), calculated from the mean abundance values across triplicate influent and effluent samples. Accompanying NMDS plots show functional divergence between influent (cyan) and effluent (red) communities.

Post-treatment shifts were also observed in the raw counts of functions related to nitrogen, phosphorus, and sulfur transformation (**Table S5: nutrient cycling**). Particularly in the agricultural ICW, the current data showed statistically significant (BH-adjusted p < 0.05) increased abundance of alkaline phosphatases (ENOG410XSIY and ENOG410YEG6, up to +9.58 log2FC) and phosphate/phosphonate transporter components (e.g., ENOG4111I6Y, up to +9.80 log2FC). In the companion animal effluent, the data showed statistically significant increases in four phosphate-selective porins O and P (ENOG410YGUH, ENOG410YHCG, ENOG410ZV9B, ENOG41101HG, up to +5.87 log2FC), phosphate/phosphonate transporters (e.g., ENOG410ZVN7, up to +8.06 log2FC), and phosphatases (e.g., ENOG4111Y3P, up to +6.12 log2FC).

### 3.6 ARG detections in influent and effluent microbial communities

Resistance genes to a vast array of antimicrobials were found in all ICW influents (**Figure 5**). ARG screening of site-specific coassemblies yielded fewer detections in the effluent compared to the corresponding influents at all sites, with reductions in the total number of distinct detected ARGs ranging from 27% for the companion animal site to >80% for the agricultural site (**Table S6**). Effluent-only detections were also observed and require validation at the sample level. At the agricultural site, >90% (118/130) of the genes detected in the influent were not detected in the treated effluent. Resistance determinants to cephalosporins, fluoroquinolones, oxazolidinones, phenicols, streptothricin, and vancomycin were removed completely or below detectable levels (**Figure 5A**). Persisting ARGs included resistance determinants to beta-lactams, aminoglycosides, macrolides, sulfonamide, tetracycline, and trimethoprim, harbored by genera such as *Pseudomonas, Legionella*, and *Acinetobacter* (**Figure 5B**). Eight genes were detected in the effluent but not in the influent, including the beta-lactamase *bla*_*OXA-29*_ belonging to *Legionella* spp. and *dfrB10*, conferring resistance to trimethoprim, belonging to unclassified members from the taxa Burkholderiales and Pseudomonadota. At the industrial site, >94% (73/77) of the genes detected in the influent were not detected in the treated effluent, including complete removal of resistance determinants to macrolides and sulfonamide. However, despite an overall lower count of ARGs detected in the effluent, 19 distinct genes were detected in the effluent but not in the influent, including the beta-lactamases *cphA5* and FOX-2 harbored by *Aeromonas*, respectively conferring resistance to carbapenems and cephalosporins, *rbpA*, conferring resistance to rifamycin, associated with *Mycobacterium*, and *vanW*, conferring resistance to vancomycin, attributed to unclassified Bacillales. The companion animal site had the largest number of detected ARGs (n=170), out of which ~50% (n=90) were removed upon treatment. In addition, 43 new ARGs were detected in the effluent, with resistance to a vast array of antibiotics including colistin, carbapenems, cephalosporins, mupirocin, streptothricin, and vancomycin. Similarly, at the municipal site, only ~45% (29/65) of the genes detected in the influent were removed in the treated effluent. Nine additional ARGs were detected in the effluent, including the carbapenemase *cphA3* belonging to *Aeromonas* spp.

**Figure 5:**
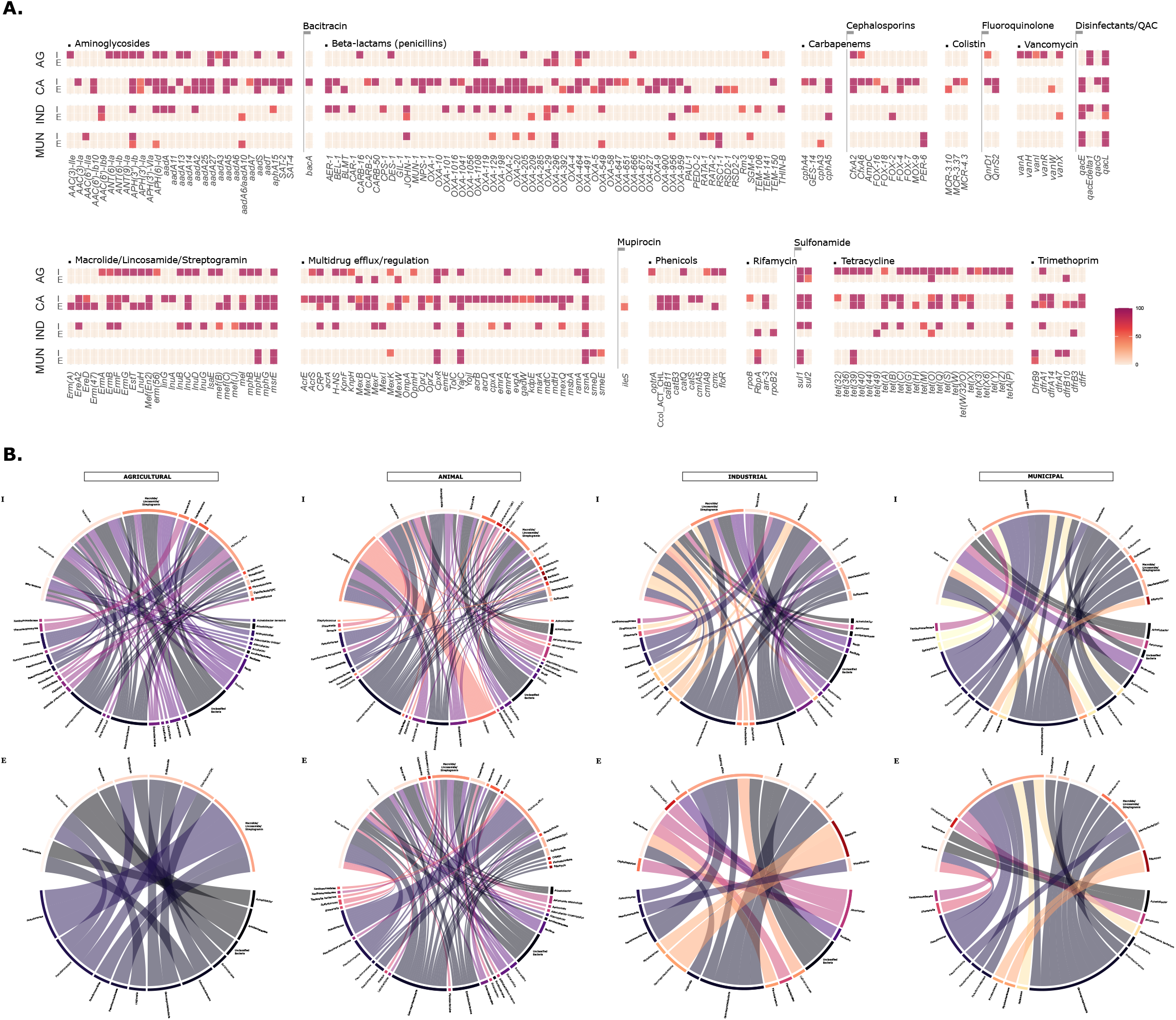
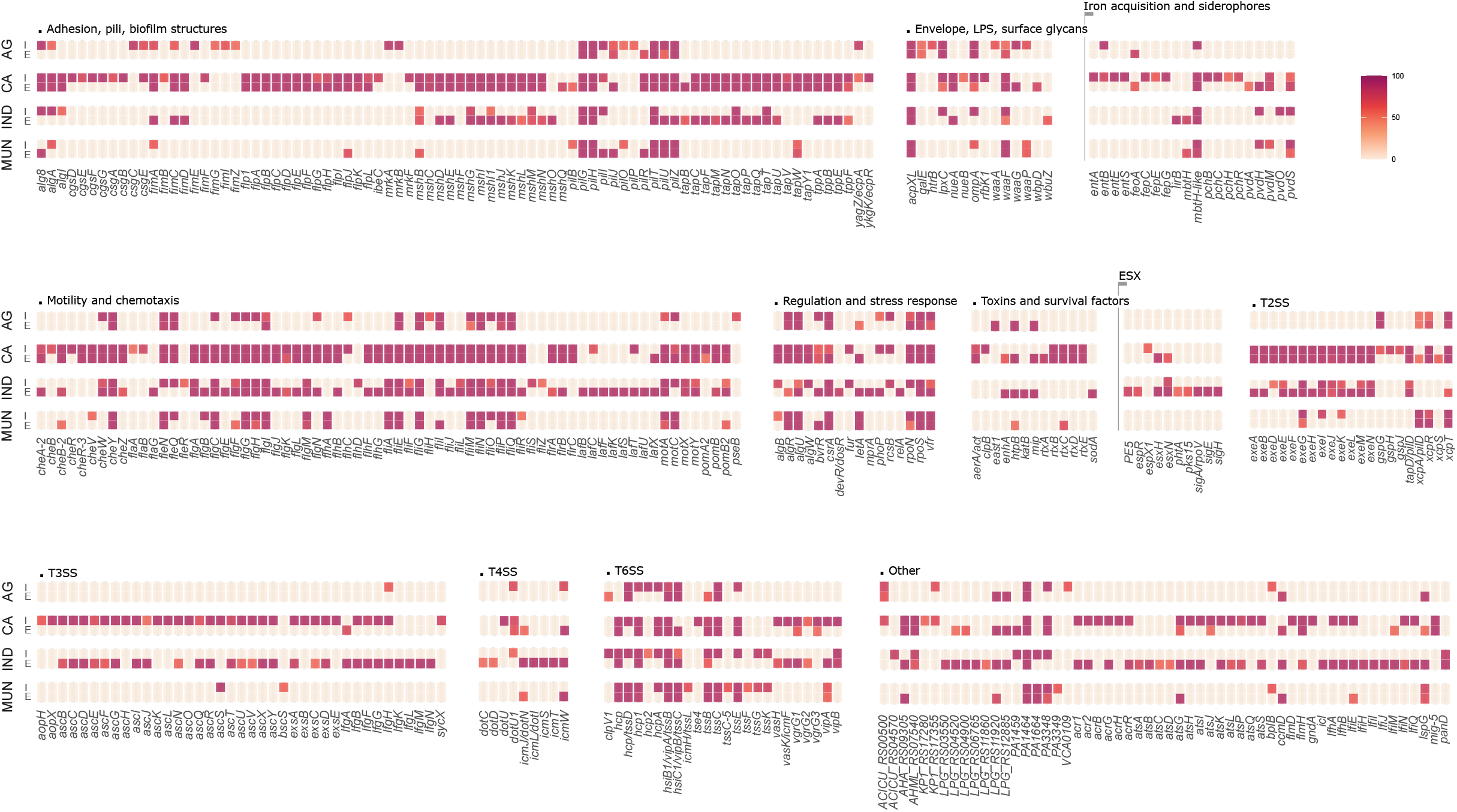
Antimicrobial resistance. (**A**) Overview of the ARGs identified in the influent (I) and effluent (E) of each sampled ICW, grouped by antimicrobial class; (**B**) Chord diagrams linking identified ARGs to their hosts, classified at the deepest possible taxonomic rank.

### 3.7 Detection of virulence and plasmid-associated markers

Screening of the coassembled metagenomic contigs against VFDB database revealed persistence across all effluents of type III (e.g., asc and exs genes) and type VI (e.g., hcp/tssD, vipA/vipB) secretion system markers associated with *Acinetobacter, Aeromonas*, and *Pseudomonas*, as well as VFs linked to *Legionella* pneumophila (*csrA, letA*, and *mip*) (**Table S7**: VFs; **Figure S1**). *Aeromonas salmonicida*-associated toxins such as *rtx* and *aerA/act* were detected in the companion animal and municipal effluents. Additionally, genes related to *Mycobacterium* persistence (*mprA, phoP*, and *sigE*) were detected in the industrial effluent. VFDB screening identified putative homologues of secretion systems and other virulence-associated genes on contigs assigned to *Acinetobacter, Aeromonas, Pseudomonas, Legionella*, and *Mycobacterium*. No ecOH O- or H-serotype loci were detected in effluent coassemblies under the applied thresholds. (**Table S7**: ecOH).

A total of 50 different replicon types were detected across all samples (**Table S7**: plasmid replicons). Particularly at the companion animal site, plasmid replicons including IncHI1A, IncHI1B, IncN, IncN5, IncC, IncP1, IncP5, IncQ1, and IncQ2 were found in both influent and effluent. IncX was detected in the influent (n=2) yet not in the effluent, while IncF was found in the influent (n=7) and only once in the effluent. Integrating the plasmid sequences and taxonomic assignment data did not show evidence of clinically relevant plasmids circulating across multiple, non-nested taxonomic groups, except for two plasmid sequences from the same replicon group (rep31) shared by Leuconostoc and Lactobacillaceae in the agricultural influent.

## 4 Discussion

Contamination from reagents, consumables, sampling, and laboratory handling is a recurring obstacle in metagenomic workflows [21,22]. Because the introduction of contaminants at any step of the workflow can distort the observed microbial community profiles, the use of negative controls is essential for robust interpretation of results. Our results reinforce this need, as sequencing the negative controls yielded over one million bacterial reads. Although confident discrimination of contaminants was complicated by the presence of shared taxa between environmental samples and negative controls, our control-informed approach allowed identification and removal of contaminant taxa from the dataset which would otherwise risk being overlooked. Additionally, the use of multiple types of negative controls implicated the cellulose nitrate membrane filters used to process the samples as the primary source of contamination (>99.9% of the negative control reads). The absence of taxa commonly associated with commercially available kits [21] suggests that the signal originated from under-characterized sources of contamination.

Nevertheless, beta diversity analyses showed separation of environmental samples from negative controls, indicating that limited contribution by contaminants to the observed microbial communities following control-informed data filtering, and overall signal reliability. Finally, close clustering of triplicates supports reproducibility and technical robustness.

Except for the municipal site, alpha diversity in treated effluents was consistently higher compared to the corresponding influents, indicating that ICWs can enhance diversity of wastewater microbial communities through treatment. As metagenomics-driven research on constructed wetlands remains globally limited, evidence on microbial dynamics is often narrow-spectrum and inconsistent, with high variability depending on wastewater type, wetland design, as well as environmental, climatic, and meteorological factors [12,23]. Limited available evidence aligns with our findings, showing a modest increase in microbial alpha diversity in municipal wastewater following treatment by a free water surface CW (observed OTUs: 614 to 691; Shannon index: 4.58 to 4.66) [23]. However, studies have also documented decreasing microbial diversity in treated effluents (Shannon index: 5.7 ± 0.2 to 4.3 ± 0.1) [12].

Influent and effluent communities separated visually in the ordinations, but the within-site tests were not significant. Together with the differential abundance data, these results suggest that such improvement in microbial diversity can occur at the expense of gut and/or human indicators and opportunistic pathogens, which are mitigated upon treatment. Of note, our results show compositional similarity of effluent samples across sites, suggesting a tendency of treated wastewater communities to converge, possibly driven by a selective biological pressure enhancing syntrophic (e.g., Syntrophorhabdaceae), fermenting (e.g., Paludibacteraceae), or nutrient cycling (e.g., *Desulfomonile*, Syntrophorhabdaceae) capabilities.

Functional characterization of wastewater metagenomes revealed that treatment significantly altered the metabolic potential of microbial communities, particularly at the agricultural and companion animal sites. A consistent decrease in functions associated with transcription and carbohydrate transport and metabolism may reflect removal of easily degradable organic matter, reduced need for rapid substrate acquisition, and overall lower microbial growth rate and metabolic activity in the effluents [24]. On the other hand, lack of significant changes at the industrial and municipal sites indicates site- and possibly source-specific treatment effects. Furthermore, our results uncovered a vast body (>80% total COGs) of unknown functions, suggesting a large reservoir of uncharacterized functional diversity in wastewater microbiomes. While this raises the possibility that key biotechnological and treatment-relevant processes may be encoded by underrepresented and poorly annotated genes, it also emphasizes the need for improved functional annotation pipelines and integrated databases.

Bidirectional changes in nitrogen- and sulfur-cycling-associated functions, together with the incomplete representation of the corresponding pathways, limited inference of coherent patterns that may indicate enhanced transformation and/or removal of these nutrients in ICW effluents. Clearer functional shifts were observed for phosphorus. In both the agricultural and companion animal ICWs, effluents were enriched in functions associated with phosphorus acquisition and mobilization, including acid and alkaline phosphatases, phosphate-selective porins, and phosphate/phosphonate transporters. Phosphate-selective porins such as OprP/OprO have previously been associated with responses to phosphate starvation [25]. Collectively, these changes are compatible with an increased capacity by the microbial community to mineralize organic phosphorus and scavenge alternative phosphorus sources in the treated effluents and may reflect selection for taxa adapted to lower bioavailable phosphorus conditions during passage through the ICW. More broadly, these observations suggest that ICW treatment may select for microbial communities with altered nutrient-acquisition and transformation potential. However, confirmation would require further in-depth, targeted nutrient chemistry measurements.

ICWs generally mitigated AMR in treated wastewater without completely eliminating ARGs from the effluents. Complete removal of ARGs is, however, not necessarily expected as these may be intrinsic to resident environmental taxa (e.g., oxacillinases in *Acinetobacter* spp.), or they may persist as extracellular DNA within biofilms and sediments. Accordingly, the detection of ARGs in ICW effluents does not itself indicate treatment failure. On the other hand, the presence of mobilizable ARGs in the environment is of greater concern, as their retention in viable bacterial hosts could provide opportunities for further dissemination through horizontal gene transfer [26]. The sulfonamide resistance gene *sul2*, usually plasmid-encoded in Gram-negative bacteria [27], and multiple aminoglycoside resistance determinants (*aadA2, aadA5, APH(6)-Id*), frequently encoded by plasmids in species such as *Enterobacter cloacae, Escherichia coli, Kluyvera georgiana, Klebsiella pneumoniae, Shigella flexneri*, and *Pseudomonas* spp. [28] were detected in association with *Enterobacter*iaceae and Gammaproteobacteria in the agricultural and companion animal effluents. Trimethoprim resistance gene *dfrB10*, found on a mega-plasmid (0.4 Mb) from *P. putida*, was also detected in association with Pseudomonadota in the agricultural effluent [29]. Plasmid- and/or transposon-mediated resistance genes to rifamycin such as *arr-3*, which is typically found in Vibrio fluvialis [30], to nucleosides (SAT-2) [31], and quinolones including QnrS2 (found in Salmonella enterica) [32], were detected on Gammaproteobacteria and *Enterobacter*ales in the companion animal effluent. The plasmid-encoded beta-lactamase *bla*_*NPS-1*_ from *Pseudomonas aeruginosa* was also detected in Pseudomonadota in the companion animal effluent [33]. Commonly plasmid-associated *mphE* and *msrE*, conferring resistance to macrolides including erythromycin [34,35], were found in Gammaproteobacteria in the companion animal and municipal effluents.

The persistence of sulfonamide, aminoglycoside, and trimethoprim resistance determinants at the agricultural ICW is compatible with the widespread use of these antimicrobial classes in Irish animal production in 2024 [36]. However, these findings should be interpreted cautiously. Because the influent wastewater consisted of stormwater runoff from the yard and roof of a dry stock farm, the residual resistome identified in the effluent likely originated from environmental bacteria and only intermittent contamination from manure and fecal material, rather than from concentrated livestock. Therefore, the detected ARGs cannot confidently be linked to patterns of agricultural antimicrobial use. The identification of plasmid-associated resistance determinants linked to *Enterobacter*ales and Gammaproteobacteria possibly represent a greater concern, as the influent wastewaters included foul waters from an animal shelter and veterinary practice. Here, the detected ARGs could have been introduced through shedding of antimicrobial-resistant bacteria in fecal material from colonized animals, as well as through wash-off of contaminated surfaces [37]. Importantly, the occurrence in treated effluent of resistance determinants to antibiotics that are not routinely used in companion animals does not imply exposure to those agents within the facility. For instance, their maintenance could result from co-selection with other linked resistance determinants which they share genomic location with [27]. Carriage by environmental bacterial taxa may provide an additional explanation. Given the single-time-point experimental design, however, these hypotheses remain speculative and cannot be assumed to represent persistent site-specific patterns.

Of further public health relevance, our results show persistence in the effluents of VFs harbored by opportunistic pathogens that are known to cause invasive human infections and contaminate water systems, such as *Aeromonas* and *Pseudomonas* spp., *L. pneumophila*, and *Mycobacterium* [38]. Furthermore, plasmid incompatibility groups commonly associated with rapid horizontal transfer and multidrug resistance persisted throughout treatment at the companion animal site. Among these, IncN, IncN5, and IncF often carry ESBLs and carbapenemase genes in *Enterobacter*iaceae [39,40], and IncHI1A, IncHI1B, and IncX are often associated with carbapenem and colistin resistance determinants in both clinical and environmental Irish isolates [41,42]. Detection of IncC plasmids in CA effluent is noteworthy as these plasmids often exhibit broad host range, possibly facilitating AMR transmission across bacterial species. IncC plasmids were previously detected in both animals and humans, suggesting a possible link with zoonotic transmission [40]. Similarly, IncP1 and IncP6 are highly relevant to wastewater surveillance, as they are often linked to AMR movement between environmental and clinical strains. Finally, small mobilizable replicons such as IncQ1 and IncQ2 detected in CA effluent often carry genes conferring resistance to sulfonamides and tetracyclines, contributing to persistent resistance reservoirs [43].

Among the 34 HQ-MAGs retrieved across the environmental samples, human or animal fecal indicators (e.g., *Agathobacter rectalis, Gemmiger qucibialis, Megamonas funiformi*) were recovered from the companion animal influent and effluent. Environmental plant- and aquatic-associated species (e.g., *Pectobacterium atrosepticum, Aquirickettsiella gammari, Shewanella oncorhynchi*) [44–46] were retrieved from the agricultural and municipal sites. Species with reported biotechnological potential were also identified in the agricultural and companion animal sites, such as *Caproicibacterium lactatifermentans* [47], previously linked with waste fermentation and caproate/medium-chain fatty acid production, and *Rhodospirillum rubrum* [48], associated with bioplastics and biohydrogen production and waste-gas valorization.

With 19 HQ-MAGs that could not be resolved to known bacterial taxa, indicating potentially novel bacterial diversity, our culture-independent investigation uncovers an uncharacterized microbial reservoir within the Irish environment which warrants further genomic and functional investigation.

## 5. Conclusions

To our knowledge, this study provides the first metagenomic characterization of microbial communities in Irish ICWs. Collectively, our findings demonstrate that ICW treatment produces substantial changes in community composition, functional potential, and AMR load, with generally increased microbial diversity accompanied by a reduction in several gut-associated taxa, opportunistic pathogens, and ARGs. Furthermore, through the analysis of the functional profile of influent and effluent microbial communities, we provide preliminary evidence of selection during ICW treatment for altered nutrient acquisition potential. Ultimately, our findings support the value of metagenomic surveillance for evaluating wastewater treatment performance.

Ongoing research on ICWs includes continuing longitudinal metagenomic-based and parallel culture-dependent monitoring of performance, building on the single-time-point findings of the present study. Future work will also enable linking predicted functional potential to microbial activity through meta-transcriptomics, and complete taxonomic and functional characterization of the identified novel bacterial species.

## Data availability

The raw sequencing data and metagenome-assembled genomes presented in this study are available on NCBI under BioProject PRJNA1456104.

## Supplementary Material Index

**Figure S1:** Overview of the virulence factors identified in the influent (I) and effluent (E) of each sampled ICW, grouped by functional class.

**Table S1:** Sample metadata. Environmental (temperature, rain, relative humidity, wind speed), water quality (sample temperature and pH), and wetland operational parameters recorded at the time of sampling.

**Table S2:** (**Domain**) Domain-level breakdown of raw read counts originating from all environmental samples (including replicates, n=24) through shotgun metagenomic sequencing. (**Negative Controls**) Domain-level breakdown of the reads originated from sequencing of the negative controls (n=10): (i) PBS filtered as the environmental samples (Filtered PBS); (ii, iii) triplicates of two cellulose nitrate membrane filters from two production batches (i.e., 251100267 and 252800617) suspended in DNA/RNA Shield without additional sample DNA; (iv) pure sterile PBS; (v) one stabilization reagent control as pure DNA/RNA Shield; (vi) one DNA extraction/sequencing negative control as DNA-free water subjected to DNA extraction and library preparation as the environmental samples. (**Genus**) Genus-level breakdown of raw read counts from both environmental samples and negative controls. (**Positive Controls**) Retrieval of the ten taxa represented in ATCC® MSA-2003™ mock community. Equal aliquots (10 µL for influent and 5 µL for effluent samples) of reconstituted ATCC® MSA-2003™ were spiked into one replicate (V=200 mL) of each sample type and used as positive controls for sequencing.

**Table S3:** Control-informed decontamination using the R package decontam. Taxa labeled as contaminant = TRUE (p.prev < 0.6) were identified as potential contaminants based on their enrichment in negative controls relative to environmental samples. “p.prev” is the prevalence-based score used to make the classification.

**Table S4:** Overview of the high-quality metagenome-assembled genomes (HQ-MAGs) retrieved from each sampled ICW. For each MAG, taxonomic classification is provided, where available, alongside key genome assembly metrics, coverage, and TPM with respect to each sample replicate.

**Table S5:** (**Overview**) Functional assignment of metagenomic contigs across all environmental samples (including replicates, n=24) and negative controls (n=8), excluding COG/ENOG IDs that were assigned to unknown functions. Only negative controls which generated functional assignments are included. (**Fig. 4B - AG; CA**) Raw data including log2 fold-changes (effluent/influent) and BH-adjusted p-values in support of **Fig. 4B**. Changes in functional feature counts were expressed as log2 fold-change (log2FC), calculated from the mean abundance values across triplicate influent and effluent samples, with a pseudo-count of 1 which was added to avoid undefined values where a feature was absent from one sample group. Positive log2FC values indicate greater abundance in the effluent, whereas a negative log2FC indicates greater relative abundance in the influent. (**Nutrient Cycling - AG; CA**) log2FC changes in functions associated with nitrogen, sulfur, and phosphorus transformation.

**Table S6:** (**Overview**) General and (**AG; CA; IND; MUN**) site-specific overview of detected antimicrobial resistance genes in the influent and effluent of each ICW. Each gene is linked to the corresponding taxonomic classification, which is classified at the deepest possible rank.

**Table S7:** Overview of detected (**VFs**) virulence factors, (**EcOH**) E. coli O- and H-serotypes, and (**plasmid replicons**) plasmid replicons in the influent and effluent of each ICW. Each identified determinant is linked to the corresponding taxonomic classification, which is classified at the deepest possible rank.

## Author contributions

**Anna Tumeo:** Methodology, Software, Formal analysis, Investigation, Data Curation, Writing - Original Draft, Writing - Review & Editing, Visualization. **Gaia Streparola:** Resources, Writing - Review & Editing. **Caolan Harrington:** Writing - Review & Editing, Funding Acquisition. **Aila Carty:** Writing - Review & Editing, Funding Acquisition. **Finola Leonard:** Writing - Review & Editing, Funding Acquisition. **Catherine Burgess:** Writing - Review & Editing, Funding Acquisition. **Dearbháile Morris:** Writing - Review & Editing, Supervision, Funding Acquisition. **Georgios Miliotis:** Conceptualization, Methodology, Writing - Review & Editing, Supervision, Project Administration, Funding Acquisition.

